# Inter-laboratory comparison of plant volatile analyses in the light of intra-specific chemodiversity

**DOI:** 10.1101/2023.02.15.528472

**Authors:** Silvia Eckert, Elisabeth J. Eilers, Ruth Jakobs, Redouan Adam Anaia, Kruthika Sen Aragam, Tanja Bloss, Moritz Popp, Rohit Sasidharan, Jörg-Peter Schnitzler, Florian Stein, Anke Steppuhn, Sybille B. Unsicker, Nicole M. van Dam, Sol Yepes, Dominik Ziaja, Caroline Müller

## Abstract

**Introduction:** Assessing intraspecific variation in plant volatile organic compounds (VOCs) involves pitfalls that may bias biological interpretation, particularly when several laboratories collaborate on joint projects. Comparative, inter-laboratory ring trials can inform on the reproducibility of such analyses.

**Objectives:** In a ring trial involving five laboratories, we investigated the reproducibility of VOC collections with polydimethylsiloxane (PDMS) and analyses by thermal desorption-gas chromatography-mass spectrometry (TD-GC-MS). As model plant we used *Tanacetum vulgare*, which shows a remarkable diversity in terpenoids, forming so-called chemotypes. We performed our ring-trial with two chemotypes to examine the sources of technical variation in plant VOC measurements during pre-analytical, analytical, and post-analytical steps.

**Methods:** Monoclonal root cuttings were generated in one laboratory and distributed to five laboratories, in which plants were grown under laboratory-specific conditions. VOCs were collected on PDMS tubes from all plants before and after a jasmonic acid (JA) treatment. Thereafter, each laboratory (donors) sent a subset of tubes to four of the other laboratories (recipients), which performed TD-GC-MS with their own established procedures.

**Results:** Chemotype-specific differences in VOC profiles were detected but with an overall high variation both across donor and recipient laboratories. JA-induced changes in VOC profiles were not reproducible. Laboratory-specific growth conditions led to phenotypic variation that affected the resulting VOC profiles.

**Conclusion:** Our ring trial shows that despite large efforts to standardise each VOC measurement step, the outcomes differed both qualitatively and quantitatively. Our results reveal sources of variation in plant VOC research and may help to avoid systematic errors in similar experiments.

## 1 Introduction

Plants produce a tremendous diversity of specialised (secondary) metabolites that differ in concentration and composition both across and within plant species (Wetzel & Whitehead, 2020). This inter- and intraspecific plant chemodiversity is particularly found in volatile organic compounds (VOCs), such as green leaf volatiles and terpenoids (Dudareva et al., 2004; Pichersky & Raguso, 2018). Constitutively produced VOCs vary among genotypes but also within individuals among different plant parts (Jakobs et al., 2019; Loreto et al., 2014; Shiojiri et al., 2021). In addition, abiotic factors, such as thermal and oxidative stress, as well as biotic factors, such as interactions with antagonists or mutualists, can induce metabolic shifts and thus alter VOC emission patterns (Dicke & Baldwin, 2010; Eberl et al., 2018; Loreto & Schnitzler, 2010). VOCs also play an important role in plant communication, for example, in pollinator attraction and direct or indirect defence against herbivores (McCormick et al., 2012; Schiestl, 2010). Intraspecific variation of emitted VOCs may affect these interactions (Aartsma et al., 2019; Kleine & Müller, 2011; Moore et al., 2014). However, assessing intraspecific variation of VOCs involves several steps that can introduce technical variation, especially when multiple laboratories collaborate on joint projects (Heil, 2014; Kallenbach et al., 2014).

The reproducibility of metabolomics approaches can be determined by using so-called ring trials (also called inter-laboratory studies or proficiency tests), in which multiple participating laboratories evaluate results obtained from a joint pool of samples (Hund et al., 2000). This approach has been applied on a variety of different analytical platforms (García et al., 2020; Izumi et al., 2019; Lin et al., 2020; Martin et al., 2015). In contrast, studies on the comparability of VOC collection and the potential sources of (technical) variation in measurements, particularly in the area of plant intraspecific chemodiversity, are underrepresented (Casadei et al., 2021; Larsen et al., 1997; Raguso & Pellmyr, 1998). VOCs are often collected using headspace sampling from entire plants or plant parts (Tholl et al., 2006). The type of adsorbent used for VOC collection, from hydrophilic to lipophilic matrices, affects the binding affinity of the individual compounds, thereby determining the detectable compound pattern, the signal intensity and sensitivity, and thus influences the results (Harper, 2000; Raguso & Pellmyr, 1998). A commonly applied adsorbent for static passive headspace VOC collections is polydimethysiloxane (PDMS; Tholl et al., 2021). After adsorption, the VOCs on the PDMS tubes are analysed via thermal desorption coupled to gas chromatography-mass spectrometry (TD-GC-MS). The tubes are of low cost and can be used for sampling in the laboratory and in the field (Kallenbach et al., 2014). Yet, it is unclear to what extent this approach might yield robust and comparable results across laboratories.

Determining VOCs includes (1) pre-analytical, (2) analytical and (3) post-analytical steps that may introduce technical variation in the obtained results (Muhamadali et al., 2020; Verpoorte et al., 2008). (1) Pre-analytical steps comprise plant cultivation and sample collection as well as sample treatment and storage (Muhamadali et al., 2020). Variation introduced during these steps may be traced back to the plant’s physiological status and the growth conditions (Niinemets et al., 2010; Tholl et al., 2006). Moreover, biological variation in VOCs may be attributed to the analysed genotype, chemotype, plant part and plant infestation status (Clancy et al., 2020; Jakobs et al., 2019; Kallenbach et al., 2014). (2) Analytical steps cover sample analysis and data collection (Muhamadali et al., 2020). Differences in analytical instruments, column length and column properties as well as measurement protocols may introduce non-biological technical variation even when the same analytical platform is used (Allwood et al., 2009). Also, relative quantities may differ due to variation in detection sensitivity, lack of peak separation and errors during peak annotation (Izumi et al., 2019). (3) Variation in post-analytical steps is caused by data processing and subsequent biological interpretation (Muhamadali et al., 2020). In order to separate these different sources of technical from the biologically interesting variation, the individual and joined effects should be investigated in inter-laboratory comparisons.

Here we aimed to quantify among-laboratory technical variation emerging from VOC sampling and analysis using common tansy, *Tanacetum vulgare* L. (Asteraceae), as a model species. This plant species displays a huge intraspecific variation of terpenoids in leaves and flowers, allowing to group individuals into distinct chemotypes (Holopainen et al., 1987; Kleine & Müller, 2011; Rohloff et al., 2004). Differences in chemotypes affect the preference and performance of different herbivores as well as their predators (Eilers et al., 2021; Jakobs & Müller, 2019; Mehrparvar et al., 2013; Wolf et al., 2012). Moreover, herbivory led to insect-specific and plant chemotype-specific induction of terpenoids and thus changes in the emission of VOCs in *T. vulgare* (Clancy et al., 2020). In general, increases in terpenoid emissions are linked to the jasmonic acid (JA) pathway, which are induced by chewing herbivores (Beckers & Spoel, 2006). Artificial JA treatments can simulate such herbivore damage (Schaller, 2008). Given that *T. vulgare* can also reproduce clonally, this species offers an ideal study system to investigate the effects of the different analytical steps in VOC sampling while keeping the genetic background constant.

To assess the reproducibility of VOC analyses via PDMS coupled with TD-GC-MS, monoclonal plants of *T. vulgare* of two distinct chemotypes were sent to five participating laboratories. Following a standardised protocol, all five laboratories collected VOCs using PDMS tubes from plants before and after JA treatment, used to mimic herbivory and enhance VOC emission. These laboratories reciprocally sent their PDMS samples to four of the five participating laboratories, in which VOCs were then measured using TD-GC-MS. We hypothesised that differences in growth among plants grown in different laboratories would introduce phenotypic variation in the number and composition of detected compounds and VOC profiles. Differences in analytical equipment and post-analytical steps were expected to introduce additional variation in the (number of) detected compounds in the recipient laboratories. Because we performed a full factorial design with two chemotypes and a JA treatment, we also tested whether the reproducibility depends on the chemotype or induction status.

## 2 Materials and methods

### 2.1 Plant growth conditions

In January 2019, seeds of two *T. vulgare* plants were collected from a population located in Bielefeld, Germany. Seeds were germinated in one laboratory (L5) on glass beads in a climate chamber (16:8 L:D, 21 °C, 70% RH). Seven-day old seedlings were transferred to pots (9 × 9 × 9 cm) containing a 1:1 mixture of steamed potting soil (Fruhstorfer Erde, Archut, Germany) and sand. Plants were re-potted to larger pots (2 L) upon growth using the same substrate, transferred to a greenhouse (21 °C; 16:8 h L:D), and fertilised weekly (modified according to Arnon & Hoagland, 1940). The terpenoid profile and thus chemotype of each plant was determined as described in Eilers et al. (2021) via GC-MS analysis from sampled leaf tissue of the second-youngest leaf. To include intra-specific variation in specialised metabolites for our inter-laboratory comparison, two individuals with distinct chemotypes were chosen. The leaf terpenoid profile of one individual was dominated by β-thujone (hereafter *mono-chemotype*) and the profile of the other was dominated by three terpenoids, i.e. (*Z*)-myroxide, santolina triene and artemisyl acetate (hereafter *mixed-chemotype*). The two individuals were reproduced monoclonally by propagating an adequate number of root cuttings to keep the genetic background within chemotype constant. The root cuttings were grown for another few weeks, before sending them to the participating laboratories.

### 2.2 Ring trial

Five laboratories (L1-L5) participated in the ring trial collecting and exchanging VOC samples from *T. vulgare* plants (hereafter *donor laboratories*), of which four of these laboratories performed the TD-GC-MS measurements of the VOC samples received (L1, L2, L4, L5; hereafter *recipient laboratories;* Fig. 1). Seven clonal plants per *T. vulgare* chemotype as well as growth substrate and pots for 14 plants plus three pots and soil for blank controls were sent via regular mail from laboratory L5 to all five participating laboratories (L1-L5) in May 2021. The delivery also contained all other material needed for performing the experiment (i.e. cups, sticks, chemicals, vials, PDMS tubes, etc). PDMS tubes (5 mm length) were prepared following Kallenbach et al. (2014; Supplementary Method S1).

**Figure 1:**
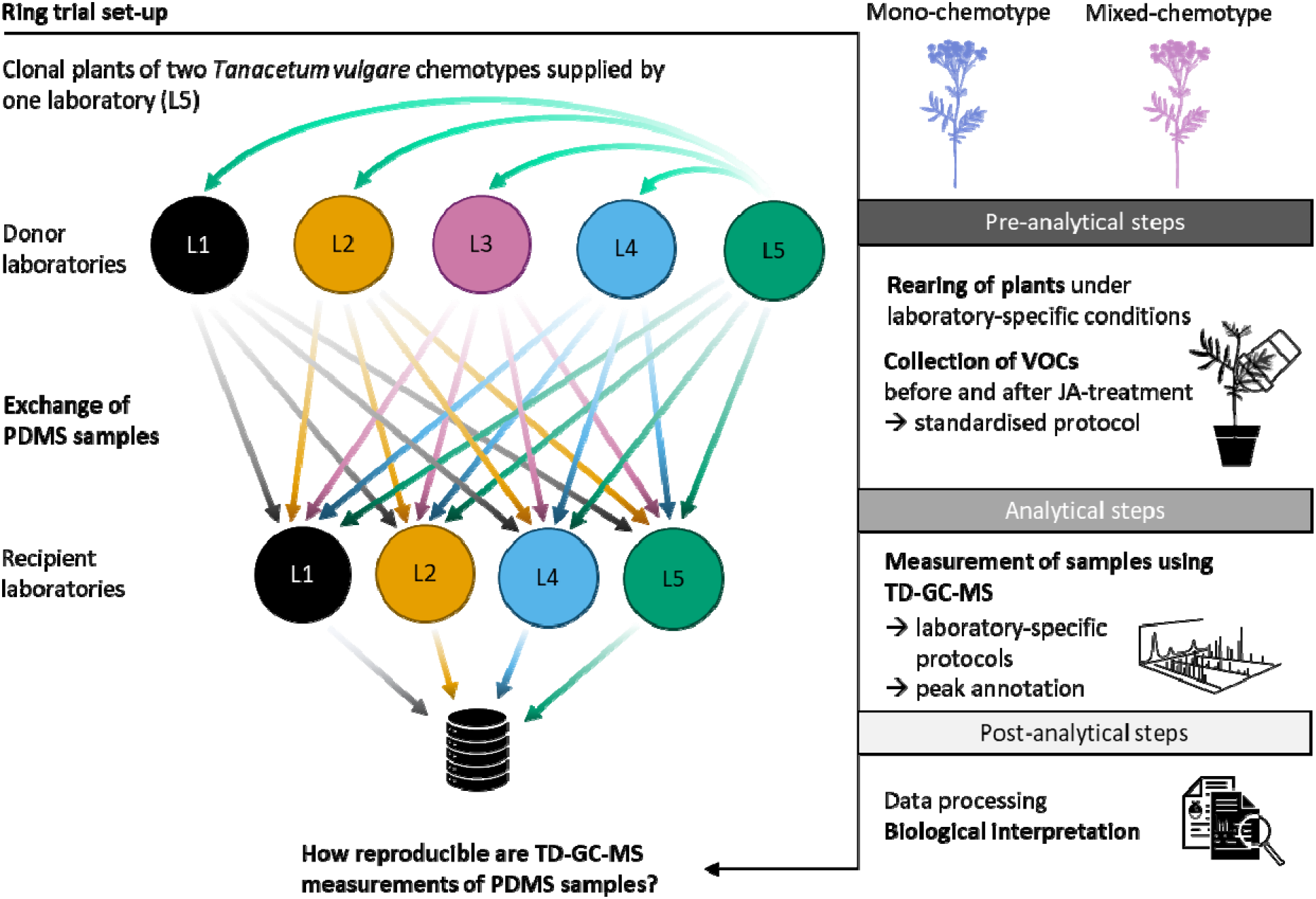
Workflow of the ring trial. Clonal plants of two *Tanacetum vulgare* chemotypes were supplied by one laboratory (L5). These plants were reared and VOCs collected with PDMS tubes before and after JA treatment using a standardised protocol in all five donor laboratories. PDMS samples were cross-exchanged with four of the participating donor laboratories (L1, L2, L4, L5; turning them into recipient laboratories) for TD-GC-MS measurements. Data of resulting VOC profiles per recipient laboratory were integrated and analysed for sources of variation.

### 2.3 Pre-analytical steps: plant growth and VOC collection before and after JA treatment

The five best developed individuals of the seven received plants per chemotype were grown in each donor laboratory under laboratory-specific conditions, i.e. either in a climate chamber (L1, L2, L4, L5, 16:8 L:D, 21 °C, 70% RH) or a greenhouse (L3; Supplementary Table S1). Plants were grown for three weeks and watered three times a week, avoiding both drought and waterlogging stress. One day before the VOC collection started, all donor laboratories determined the plant height and the number of expanded leaves as phenotypic parameters. VOC collection was performed by each donor laboratory according to a strict protocol (for details see Supplementary Method S1 and Fig. S1) as follows: 4 h after the onset of the photoperiod, the youngest fully developed leaf per plant was enclosed in a 600 mL polyethylene terephthalate (PET) cup (Wimex, Náchod, Czech Republic) by inserting it through a hole made in the bottom of the cup. The cup was fixed in position with a balloon stick. Plants were allowed to recover from handling stress for one day, as mechanical damage after handling can lead to higher VOC emission rates (Loreto et al., 2000).

Four hours after the onset of the photoperiod of the following day, the VOC collection was initiated. Ten μL of 100 ng μL^-1^ 1-bromodecane (Sigma-Aldrich Chemie GmbH, Taufkirchen, Germany) solved in *n*-heptane (GC-MS grade; CHEMSOLUTE, Th. Geyer GmbH & Co. KG, Renningen, Germany) was applied on a filter paper (1 cm^2^) and allowed to evaporate for a few minutes. Then, the paper was inserted through the opening of the cup and placed onto the leaf, serving as internal standard. Next, twelve PDMS tubes were inserted per cup using cleaned forceps, ideally neither touching each other nor the leaf. To assess the volatile background, blank samples using empty cups were prepared (hereafter blank controls). A balloon stick was used to hold the empty cup above a pot containing substrate only. As for the leaf sampling, one filter paper with internal standard and twelve PDMS tubes were inserted per cup. The relative air humidity, light intensity and air temperature were measured once during the VOC collection. After six hours, the PDMS tubes were removed from the cups without touching the leaves and equally divided into six 1.5 mL glass vials, resulting in two PDMS tubes per vial. The glass vials were sealed with polytetrafluoroethylene (PTFE) tape and stored at −20 °C until they were mailed to the four recipient laboratories or analysed in the own lab. The filter paper with the 1-bromodecane was discarded.

One day after the first VOC collection and four hours after onset of light, a herbivore damage was simulated by treating all plants with 10 mL of a 0.005 % JA solution [JA diluted in double de-ionised water containing 0.1% (v:v) Triton X-100 as surfactant (Sigma-Aldrich)]. The JA solution was injected in three portions into the substrate around the stem of each plant as well as into the substrate of the control pots. In some laboratories, it was noted that the JA did not go in solution very well. One day after the JA application, again a new filter paper freshly treated with 1-bromodecane and twelve fresh PDMS tubes were inserted into each cup. For nine plant samples in L3, no sufficient 1-bromodecane was left and could thus not be added. This was also the case for twelve blank samples in L1, four blank samples in L2, and nine blank samples in L3. Environmental conditions were measured again, as described above. After six hours of sampling, PDMS tubes were removed, distributed in separate glass vials sealed with PTFE tape, with two tubes per vial, and vials frozen at −20 °C. The plastic cup was removed and the leaf, which had been VOC-sampled, was cut at the base of the petiole. The fresh weight of this leaf was determined. Finally, the entire aboveground biomass was harvested and weighed.

From each sampled plant, one vial with PDMS tubes collected before and one with tubes collected after the JA treatment as well as vials with PDMS tubes from the respective blank controls were sent to each of the recipient laboratories (L1, L2, L4, L5). After shipping without additional cooling, samples were again stored at 20 °C for at least three weeks until analyses. For unexplained reasons, samples sent by L3 did not arrive at L1.

### 2.4 Analytical steps: TD-GC-MS analyses

Recipient laboratories analysed the received PDMS tubes using a protocol modified after Kallenbach et al. (2014), adjusted to their TD-GC-MS instrument (Table 1). Samples were taken out of the freezer 24 h before TD-GC-MS analysis. Per plant and treatment level, one PDMS tube was analysed, with all samples measured in a randomised order. Each batch of measurements was accompanied by measurements of a PDMS tube with a mixture of alkanes (C7–C40, Sigma-Aldrich) to calculate retention indices (RI; Adams, 2007; van Den Dool & Kratz, 1963). An empty TD-glass tube (without any PDMS tube) was measured as background control. Additionally, standards of camphor, β-caryophyllene, *p*-cymene, α-pinene (all Sigma-Aldrich) and (*Z*)-hex-3-enyl-acetate (Thermo Fisher, Kandel, Germany) were applied directly in TD-glass tubes and measured in a concentration series (20-100 ng μL^-1^; Supplementary Fig. S2).

**Table 1:** Overview of TD-GC-MS instruments and settings for analyses of PDMS tubes used in the different recipient laboratories.

|  | L1 | L2 | L4 | L5 |
| --- | --- | --- | --- | --- |
| **Instrument** | Shimadzu TD-20<br>GC-MS-QP2010Ultra | Agilent GCMS<br>5975C with 7890<br>GC and Gerstel<br>Maestro | Shimadzu TD-30,<br>GCMS<br>QP2020NX | Shimadzu TD-20, GC<br>2010 Plus QP2020 |
| **Column** | ZB-5 (30 m ×<br>0.25 mm, 0.25<br>µm film<br>thickness) | J&W 122-5562G_1 – DB-5MS + 10 m DG<br>(69 m × 0.25<br>mm, 0.25 µm<br>film thickness) | Rtx-5MS (30 m ×<br>0.25 mm, 0.25<br>µm film<br>thickness) | VF5-MS (30 m ×<br>0.25 mm, 0.25 µm<br>film thickness) |
| **TD settings:** |  |  |  |  |
| GC interface (transfer line) | 230 °C | 230 °C | 250 °C | 250 °C |
| Sampling (flow rate and<br>temperature) | 60 mL min<sup>-1</sup>,<br>230 °C | 100 mL min<sup>-1</sup>,<br>ramp from -20°C<br>to 270°C | 60 mL min<sup>-1</sup> for<br>8 min at 210 °C | 60 mL min<sup>-1</sup> for 8<br>min at 210 °C |
| Cryotrap cooling<br>temperature | -17 °C | -20 °C | -15 °C | -15 °C |
| Cryotrap heating<br>temperature | 230 °C, hold for 5<br>min | 12 °C min<sup>-1</sup> to<br>270 °C, hold for 2<br>min | 230 °C, hold for<br>10 min | 230 °C, hold for 10<br>min |
| **GC settings:** |  |  |  |  |
| Carrier gas | helium | helium | helium | helium |
| Flow control mode | Linear velocity | Constant flow | Linear velocity | Linear velocity |
| Total flow | 13.7 mL min<sup>-1</sup> | 104 mL min<sup>-1</sup> | 13.8 mL min<sup>-1</sup> | 13.8 mL min<sup>-1</sup> |
| Column flow | 1.46 mL min<sup>-1</sup> | 1 mL min<sup>-1</sup> | 1.46 mL min<sup>-1</sup> | 1.46 mL min<sup>-1</sup> |
| Purge flow | 5 mL min<sup>-1</sup> | 3 mL min<sup>-1</sup> | 5 mL min<sup>-1</sup> | 5 mL min<sup>-1</sup> |
| Pressure | 85.7 kPa | 127.9 kPa | 114.7 kPa | 114.7 kPa |
| Split ratio | 5 | splitless | 5 | 5 |
| Oven program | 50 °C for 5 min,<br>ramp to 250 °C<br>with 10 °C min<sup>-1</sup>, | 40 °C; 50 °C min<sup>-1</sup><br>until 50°C, hold 5<br>min; 10 °C min<sup>-1</sup> | 50 °C for 5 min,<br>ramp to 250 °C<br>with 10 °C min<sup>-1</sup>, | 50 °C for 5 min,<br>ramp to 250 °C with<br>10 °C min<sup>-1</sup>, ramp to |
|  | ramp to 280 °C<br>with 30 °C min<sup>-1</sup>,<br>hold for 2 min | until 250 °C; 30<br>°C until 300 °C,<br>hold 3 min | hold for 2 min | 280 °C with 30 °C<br>min<sup>-1</sup>, hold for 2<br>min |
| Total duration of GC run | 28 min | 31 min | 27 min | 28 min |
| **MS settings:** |  |  |  |  |
| Transfer line temperature | 250 °C | 250 °C | 250 °C | 250 °C |
| Ion source temperature | 230 °C | 230 °C | 230 °C | 230 °C |
| Scan range | Full scan from 33<br>to 400 *m/z* | Full scan from 35<br>to 350 *m/z* | Full scan from 30<br>to 400 *m/z* | Full scan from 30 to<br>400 *m/z* |
| Offset starting time for MS<br>measurements | 0 min | 9.8 min | 0 min | 7.0 min |

### 2.5 Post-analytical steps: compound annotation and integration

Recipient laboratories used their laboratory-specific infrastructure to annotate compounds from the obtained TD-GC-MS chromatograms. Compounds were annotated comparing the RI to those published by Adams (2007) and available libraries (Supplementary Table S2). To facilitate peak annotation, one recipient laboratory (L5) distributed a reference list with all previously identified compounds in *T. vulgare* samples. Compounds were semi-quantified based on the extracted ion chromatogram (EIC) of the corresponding peak spectrum. Obtained peak areas were normalised according to the fresh weight of the respective VOC-sampled *T. vulgare* leaf.

### 2.6 Statistical analyses

All data were analysed with the statistical software R v4.1.2 (R Core Team, 2022). To test whether laboratory-specific environmental conditions affected phenotypic variation of the plants, each of the four measured traits (i.e. plant height, number of leaves, fresh aboveground biomass and fresh mass of VOC-sampled leaf) was compared using Type III (in case of significant interactions) and Type II (in case of significant main effects) two-way analysis of variance (ANOVA). Donor laboratory, chemotype and their interaction were used as explanatory factors and models calculated using the *lm* function from the stats v4.2.1 package and *Anova* function from the car v3.1-0 package (Fox et al., 2019). Post-hoc testing was applied using Tukey’s range test with the *HSD.test* function from the agricolae v1.3-5 package (Mendiburu, 2021). Assumptions of linear models were checked using the *check_model* function from the performance v0.9.1 package (Lüdecke et al., 2021). In the case of non-normality of residuals, response variables were transformed using ordered quantile normalisation (the number of fully unfolded leaves; Peterson & Cavanaugh, 2019) and Box-Cox transformation (fresh weight of VOC-sampled leaf; Box & Cox, 1964).

To assess whether the number of samples in which 1-bromodecane was detected by recipient laboratories differed between donor laboratories, permutation-based analysis of variance (LM_Perm_) was performed using the *aovp* function from the LMPerm v2.1.0 package (Wheeler & Torchiano, 2016) with donor laboratory as explanatory factor. We also used chemotype as an additional explanatory factor to assess whether the recovery of 1-bromodecane in a sample is influenced by the specific emitted plant volatiles during handling. In the model, the number of samples in which 1-bromodecane was detected was applied as a relative response (in percentage per recipient laboratory) and normalised using ordered quantile normalisation. To test whether the use of 1-bromodecane as an internal standard might yield reproducible results when applying the PDMS approach, intraclass correlation coefficients (ICC) among recipient laboratories were calculated separately for each chemotype but also for blank samples. For this, the *icc* function from the irr v0.84.1 package (Gamer et al., 2019) was used on single-value rating, i.e. normalised peak area, with both subjects and raters, i.e. recipient laboratories, defined as randomly chosen to assess interrater agreement. The distribution of 1-bromodecane (normalised peak area) found in samples before and after JA treatment did not significantly differ (Mann Whitney *U* test; mono-chemotype: U = 4181.5, *p* = 0.583; mixed-chemotype: U = 4143, *p* = 0.795) and samples were therefore pooled for both LM_Perm_ and the ICC.

Separate Venn diagrams per *T. vulgare* chemotype were plotted to determine the degree of overlap between the VOC profiles measured by the different recipient laboratories (Venn, 1881). Only VOCs with a RI > 933 were included (Supplementary Table S2). To test whether handling in the donor laboratories affected the variation in detected VOCs per sample, the number of peaks per sample was analysed as response variable in longitudinal linear mixed-effects models (LMM) using the chemotype as well as donor laboratory and their interaction as fixed effects. Additionally, recipient laboratory, treatment and plant individual nested within recipient laboratory were used as random effects. A similar LMM was applied to test whether measurement in the recipient laboratories affected the number of detected VOCs, switching both donor laboratory (here random effect) and recipient laboratory (here fixed effect). The importance of each fixed factor per LMM was assessed using likelihood-ratio tests where the original model was compared with a model without interaction and the non-interaction model was compared to separate models lacking each of the main factors. For all further statistical analyses, only six jointly detected volatile terpenoids (borneol, camphor, *p*-cymene, eucalyptol, α-pinene and β-thujone) were compared between donor laboratories and recipient laboratories, respectively.

To capture the variability of the obtained VOC profiles by the participating donor and recipient laboratories, respectively, non-parametric multidimensional scaling (NMDS) was performed, using the Kulczynski distance on square root-transformed and Wisconsin double-standardised data in the *metaMDS* function of the vegan package. To facilitate the convergence of the NMDS solution due to zero values, a very small number was added (1e^-10^) before data transformation. The distance matrix was applied to a multivariate analysis of dispersion (betadisper) using the *betadisper* function in the vegan v2.6-2 package (Oksanen et al., 2022) to check whether the variability between samples differed in a laboratory-specific and chemotype-specific manner. The betadisper analysis was followed by a non-parametric multivariate analysis of variance (npMANOVA) to test for chemotype-specific variation in the VOC profiles using the *adonis2* function in the vegan package. In npMANOVA with chemotype as explanatory variable, permutations were constrained to recipient laboratories nested within donor laboratories. For both betadisper analysis and npMANOVA as well as the stacked barplots, only data (relative amount in percentage) from the untreated plants were used to test for reproducibility of the PDMS approach without the confounding effect of defence induction. VOC profiles of untreated leaves were visualised using stacked barplots split by chemotype, donor laboratory and recipient laboratory, respectively.

To assess whether the resulting VOC profile per sample was affected by early-stage phenotypic variation of plants, a distance-based redundancy analysis (dbRDA) was applied using the distance matrix from NMDS and the *capscale* function from the vegan package. To avoid collinearity of the explanatory variables, from the four plant traits measured one representative trait (plant height) was selected from a correlation matrix using Pearson correlation, as it showed the highest correlation. Significance in dbRDA was assessed using permutation tests based on 9999 permutations. To test whether the changes in VOC profiles according to JA treatment are consistent, the log_2_-fold change (log_2_FC) was calculated per donor laboratory on each of the compounds jointly detected by the recipient laboratories. Log_2_FC was calculated for each recipient laboratory within each donor laboratory using a modified version of the *omu_summary* function from the omu v1.0.7 package (Tiffany & Bäumler, 2019), and the median-averaged log_2_FC was used for comparison. To avoid infinite numbers in the calculation of log_2_FC values, a constant was added that was two orders of magnitude lower than the peak area of the smallest compound detected per recipient laboratory (ten for L1, L2, and L5; one for L4), placing zero-value peak areas within the background noise. Variation in log_2_FC values was analysed based on individual log_2_FC values using longitudinal linear mixed-effects models with donor laboratory, VOC and their interaction as fixed effects, and recipient laboratory as well as plant individuals nested within recipient laboratory as random effect. Models were applied separately for each chemotype and log_2_FC values were normalised using ordered quantile normalisation. The significance of model terms was analysed with likelihood-ratio tests by comparing the original model to models reduced for each model term. Univariate data were visualised with either a point-range plot using the arithmetic average and standard deviation (i.e. for number of detected 1-bromodecane peaks per recipient laboratory), box-and whisker plots or a heatmap (i.e. the effect of JA induction).

## 3 Results

### 3.1 Pre-analytical steps: variation in plant size and reproducibility of recovery of internal standard

The number of leaves per plant, measured one day before VOC collection in untreated plants, and plant height of the two chemotypes varied significantly depending on the donor laboratory (Fig. 2A, B; Supplementary Table S3 and S4). Plants of the mixed-chemotype were smallest when grown in laboratory L1, while all plants grown in L1 had about four times more leaves than plants grown in the other laboratories. Fresh aboveground biomass differed significantly among donor laboratories, being highest in plants grown in L4 and lowest in those grown in L2 (Fig. 2C; Supplementary Table S4). Shoots of the mono-chemotype were significantly heavier than shoots of the mixed-chemotype. The fresh weights of the VOC-sampled leaves differed significantly between chemotypes depending on the donor laboratory involved, being particularly low in the mono-chemotype of L2 and the mixed-chemotype in L4 (Fig. 2D; Supplementary Table S3).

**Figure 2:**
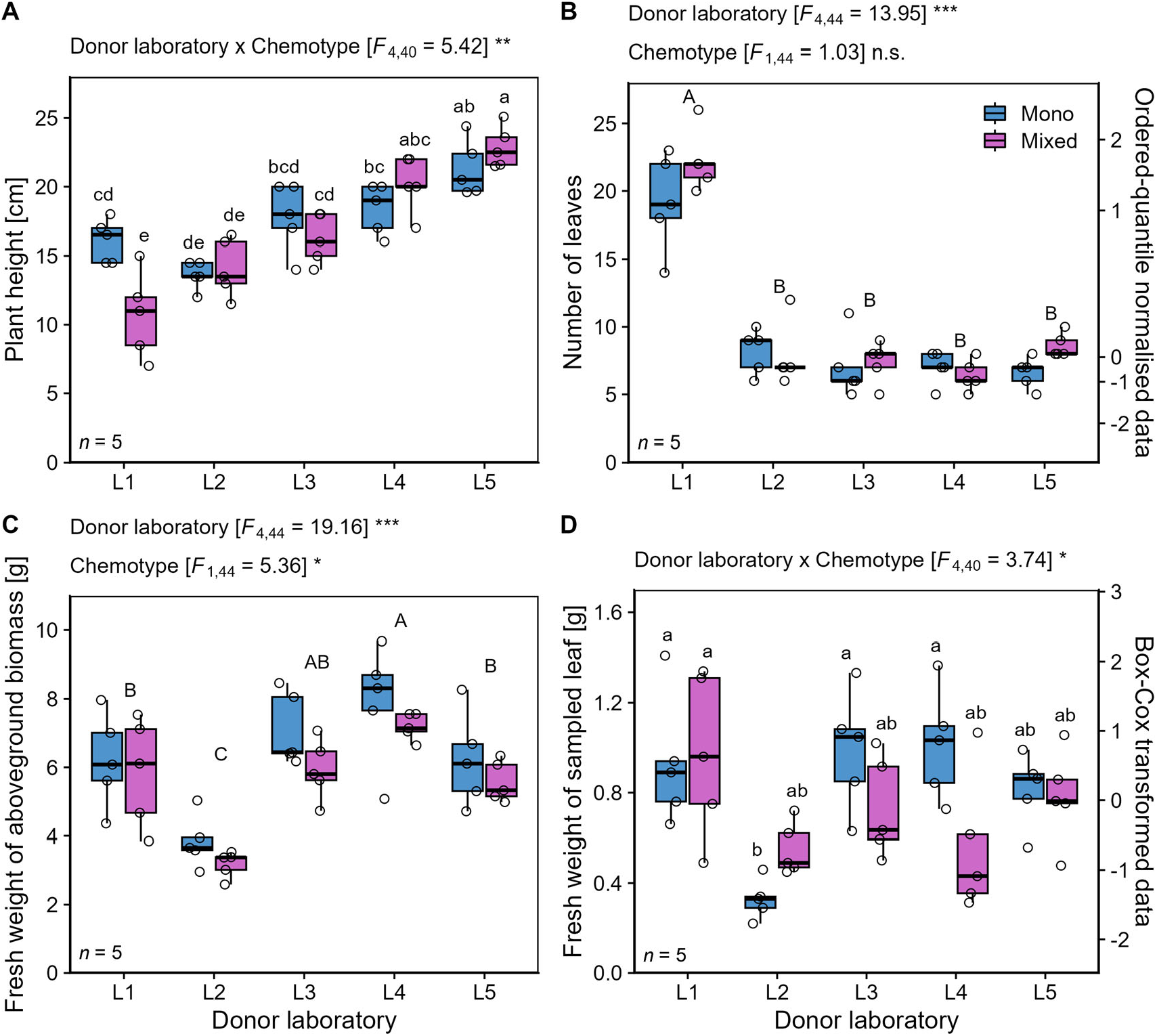
Phenotypic variation in *Tanacetum vulgare* plants of two chemotypes grown in five donor laboratories (L1-L5). The traits **(A)** plant height [cm] and **(B)** number of leaves were measured one day before VOC sampling, while the fresh weight of **(C)** aboveground biomass (shoots) and **(D)** the VOC-sampled leaf were harvested at the end of the experiment (three days later). Abbreviations: Asterisks denote significance levels in two-way ANOVA at *p* < 0.001 (***), *p* < 0.01 (**) and *p* < 0.05 (*); n.s. – not significant. Results from post-hoc testing using Tukey’s range test are indicated with capital letters for significant pairwise differences between donor laboratories and lowercase letters for significant pairwise differences in the interaction of chemotypes and donor laboratories. Number of replicates per chemotype and donor laboratory: *n* = 5. Data transformations are displayed on a second y-axis. Data are presented as box-and-whisker plots with interquartile ranges (IQR, boxes) including medians (horizontal thick lines), whiskers (extending to the most extreme data points with maximum 1.5 times the IQR) and raw data points (open circles).

Overall, 1-bromodecane peaks were found in 45% of all samples analysed (166 out of 371 samples). The number of samples, in which the internal standard was detected, did only marginally differ among laboratories (*p* = 0.120) and was not affected by chemotype (Fig. 3A; Supplementary Table S5). The quantity of the internal standard was not reliably determined between recipient laboratories for the mono-chemotype (ICC close to 0; Fig. 3B) but was significantly reproduced for the mixed chemotype, albeit with poor agreement (ICC < 0.5) according to Koo & Li (2016). The quantity of 1-bromodecane was also not reliably determined in blank samples (Supplementary Fig. S3).

**Figure 3:**
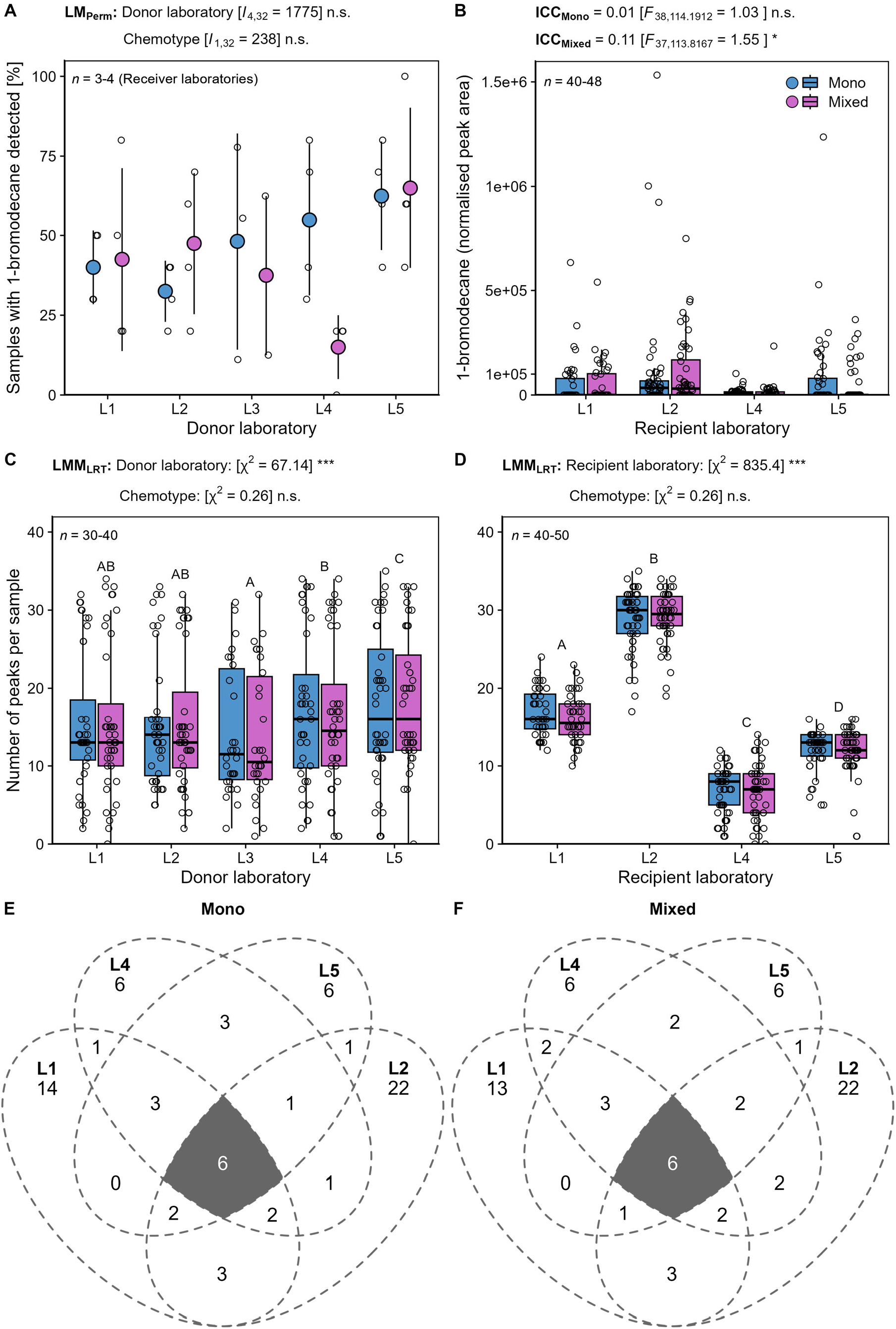
Variation in detected peaks for both internal standard (1-bromodecane) and VOCs per chemotype of *Tanacetum vulgare* and laboratory. **(A)** Number of samples per recipient laboratory, in which 1-bromodecane was detected per donor laboratory and **(B)** the normalized peak area of 1-bromodecane per recipient laboratory is shown. The number of detected peaks per sample and chemotype is given separately for **(C)** donor laboratories and **(D)** recipient laboratories. The total numbers of VOCs detected by TD-GC-MS from PDMS sampling overlapping (grey area) or found exclusively in the participating recipient laboratories are presented for **(E)** the mono-chemotype and **(F)** the mixed-chemotype. Data are shown irrespective of treatment levels as **(A)** point-range plot based on averages and standard deviations, **(B, C, D)** box-and-whisker plots, presenting the median within 50 % of data in boxes and 1.5-times the inter-quartile range as whiskers and raw data points (open circles; **A-D**) and **(E, F)** Venn diagrams. Abbreviations: Asterisks denote significance levels in permutation-based linear models (LM_Perm_), intra-class correlation coefficients (ICC), and likelihood-ratio tests (LRT) from linear repeated-measures mixed-effects models (LMM) at *p* < 0.001 (***)and *p* < 0.05 (*); n.s. – not significant. Please note that, for better visualisation, one outlier at 3.11 e^+6^ was omitted in **(B)**.

### 3.2 Pre-analytical and analytical steps: variation in number of compounds detected across all laboratories

Based on EIC-mode analyses, the number of peaks detected per sample differed significantly among both samples sent by different donor and recipient laboratories, while it was not affected by chemotype (Fig. 3C, D; Supplementary Table S6 and S7). Among the participating recipient laboratories, the lowest number of VOCs was detected in L4 (maximum of 23 and 25 peaks, respectively) and the highest number of VOCs was detected in L2 (38 and 39 peaks, respectively; Fig. 3C, D). In total, 72 VOCs were detected in plants of both chemotypes with six of these VOCs jointly detected by all recipient laboratories (Fig. 3E, F; borneol, camphor, *p*-cymene, eucalyptol, α-pinene and β-thujone, respectively; Supplementary Table S2). The VOC profile of these jointly detected VOCs was significantly affected by the plant height of individuals reared at the different donor laboratories (dbRDA: *F*_1,188_ = 5.80, *p* < 0.001) pointing to an impact of phenotypic variation generated during pre-analytical steps on later analytical outcomes.

### 3.3 Post-analytical steps: commonly detected compounds and variation in VOC profiles among laboratories and due to JA treatment

The composition of the six jointly detected terpenoids per recipient lab (borneol, camphor, *p*-cymene, eucalyptol, α-pinene and β-thujone) was dominated by β-thujone in the mono-chemotype (15.8-93.2%), although the abundance varied especially for *p*-cymene (2e^-11^-46.7%) and α-pinene (3.1 – 45.7%; Fig. 4A). The VOC profiles of the mixed-chemotype were not dominated by a certain terpenoid, however, β-thujone dominated in the VOC profiles of L5 (11.7 – 63.1%). The degree of variability of detected VOC profiles varied significantly for both donor laboratories and recipient laboratories, but not between the two chemotypes (betadisper analysis; Fig. 4B, C). In addition, the relative proportions of these six VOCs in the blend differed significantly between the two chemotypes (Fig. 4C). For both chemotypes, all six VOCs responded to the JA treatment, but bi-directionally and inconsistently between donor laboratories (Fig. 5). Eucalyptol showed the highest increase in log_2_FC in L2 and L3 consistently in both chemotypes, whereas β-thujone showed the strongest decrease in log_2_FC in L1 for the mono-chemotype and L5 for the mixed-chemotype. Little to no response was found for the mono-chemotype for borneol and *p*-cymene for L5 (Fig. 5A). For the mixed-chemotype, little to no response was detected in borneol and camphor for L5 and L3, respectively. A chemotype-specific compound profile could not be reproduced across donor laboratories, as log_2_FC values varied significantly among compounds depending on the donor laboratory involved (Fig. 5A, B; Supplementary Table S8).

**Figure 4:**
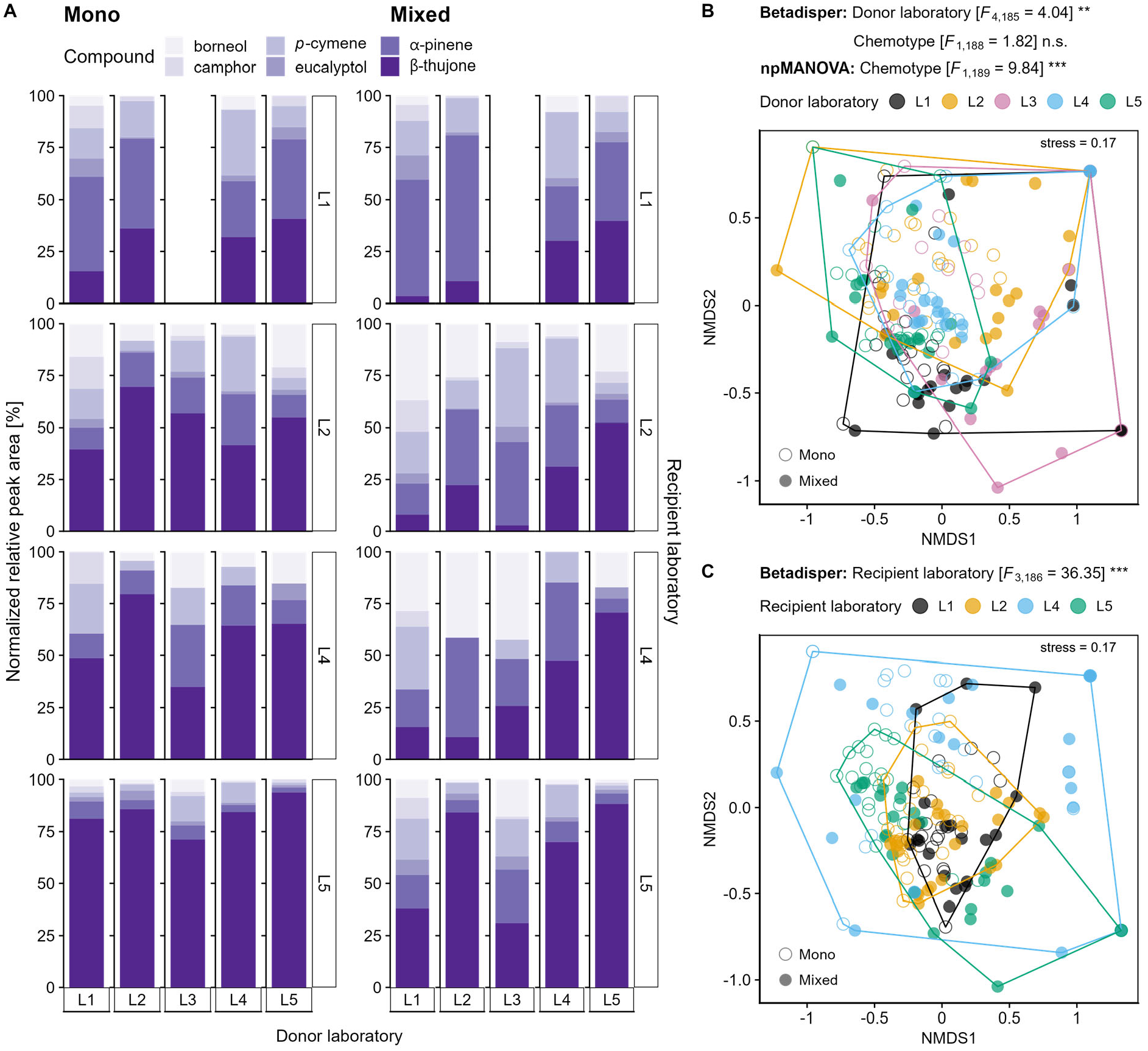
Variation in VOC profiles of two chemotypes of *Tanacetum vulgare* chemotypes sampled and analysed in different laboratories before jasmonic acid-treatment. The profiles of six jointly detected VOCs collected via PDMS sampling are shown as comparison between **(A)** chemotypes (filled circles: mono-chemotype, open circles: mixed-chemotype), **(B)** donor laboratories (performing pre-analytical steps), and **(C)** recipient laboratories (responsible for analytical steps). Data are shown as **(A)** as percentage of normalised peak area (average of *n* = 5 samples) and **(B, C)** NMDS plots based on percentage of normalised peak areas that were squared and standardised using Wisconsin double standardisation. Abbreviations: Asterisks denote significant differences in betadisper analysis and two-way npMANOVA at *p* < 0.001 (***) and *p* < 0.01 (**).

**Figure 5:**
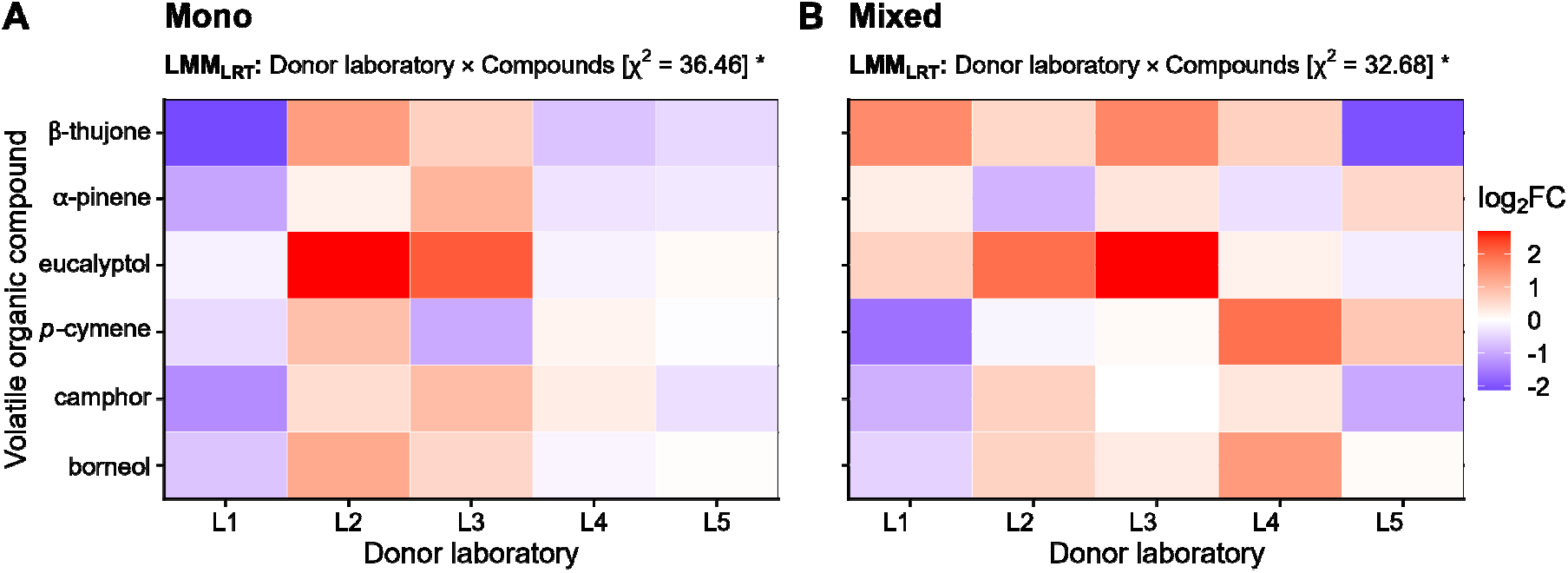
Effect of jasmonic acid (JA) treatment on VOC profiles. Differences between VOC amounts in the **(A)** mono- and **(B)** mixed-chemotype of *Tanacetum vulgare* (each with a sample size of *n* = 5) before and after treatment with JA plotted as median log2-fold changes (log_2_FC) across recipient laboratories. Log_2_FC values lower or higher than zero denote a decrease or increase of the VOC concentration after JA treatment, respectively. Abbreviations: Asterisks denote significance levels in likelihood-ratio tests from linear repeated-measures mixed-effects models (LMM_LRT_) at *p* < 0.05 (*); n.s. – not significant.

## 4 Discussion

In recent years, the reproducibility of scientific results has been called into question (Ioannidis, 2005) and this problem has also been acknowledged in metabolomics studies (Ghosh et al., 2021; Stavarache et al., 2022). Using a comparative inter-laboratory ring trial, we aimed at identifying sources of variation that may influence the reproducibility of obtained results on plant VOC profiles when applying the PDMS approach coupled with TD-GC-MS analysis in collaborative projects. Using clones of two chemotypes of *T. vulgare*, we found that laboratory-specific conditions of the donor laboratories as well as TD-GC-MS equipment on which VOCs were analysed in the recipient laboratories influenced the recovery of VOC profiles in both *T. vulgare* chemotypes. In the following, we discuss potential sources of variation at the different experimental steps, which can impede the reproducibility of results from PDMS sampling measured with TD-GC-MS.

### 4.1 Variation introduced by pre-analytical steps

Despite growing clonal material of just two *T. vulgare* source plants in all laboratories, the plants showed individual differences in size before shipping and when grown under laboratory-specific environmental conditions in the different laboratories. Particularly the light conditions were quite distinct among the donor laboratories. In addition, plants may have been exposed to different conditions during mailing to all donor laboratories. High phenotypic plasticity as a response to environmental cues has been found in various clonal plant species and is an important part of biological variation (Fazlioglu & Bonser, 2016; Liu et al., 2016; Price & Marshall, 1999). Moreover, propagating plants through cuttings has been found to be epimutagenic, inducing changes in the epigenome associated with the abiotic conditions experienced by the propagules (Lloyd & Lister, 2022). Additionally, genotype-by-environment interactions can enhance intraspecific variation in functional traits (Barker et al., 2019; Des Marais et al., 2013). Such interaction may have led to the large variation in plant height and the weights of the VOC-sampled leaves in the *T. vulgare* plants grown under slightly varying laboratory-specific environmental conditions in the present experiment. The variation in plant height, which correlated with the other growth-related traits measured, could indeed explain the variation in VOC profiles detected as indicated by the dbRDA, highlighting the importance of variation introduced in pre-analytical steps in ring trials. Intra-clonal variation has also been found, for example, in vegetatively propagated plants of *Pelargonium* sp. (Geraniaceae) for the content and composition of essential oils, but not for morphological traits (Kulkarni et al., 1997). In contrast, in another plant species, *Tropaeolum majus* (Tropaeoloaceae), variation in chemical defence traits such as glucosinolates seemed to vary only little within clones but levels were expected to increase under changing environmental conditions (Kleinwächter et al., 2008). Plant species showing a high intra-clonal variation in functional and defence traits may be particularly challenging when aiming to test for reproducibility of chemical analyses. Furthermore, the preparation of biological material, e.g. the way how treatments are applied as well as the handling of collected metabolite samples, might affect the outcome of later analyses (Muhamadali et al., 2020). We tried to reduce such factors, which could introduce variation, to a large extent. For example, touching of plants can lead to destruction of the extracellular terpenoid-containing glandular trichomes (Lange and Srivifya, 2019) and induce their VOC emission, as found in *T. vulgare* plants when enclosing them into cuvettes (Clancy et al., 2020). In our experiment, plants had to be touched to apply the cups in which PDMS tubes were inserted. However, the fact that we enclosed the leaves already one day before the VOC collection into the cups and entered the PDMS tubes without having to touch the plants might have prevented or at least reduced a potential effect of different VOC release.

In addition, laboratory-specific micro-climatic conditions might influence the quantitative and qualitative composition of emitted VOCs. For example, the emission of mono- and sesquiterpenoids in *Quercus coccifera* (Fagaceae) was found to respond to changes in light and temperature conditions even within short term (Staudt & Lhoutellier, 2011). In contrast, recovery of VOCs sampled by PDMS tubes has been proven to be quite robust towards differences in incubation temperature, with only modest effects on recovery between 4 °C to 50 °C, but critical effects were found with respect to sampling time (Kallenbach et al., 2014). Therefore, sampling time was highly standardised in the present experiment, with six hours across all laboratories. Also belowground conditions, such as soil humidity and fertilisation rate, might affect the quantity and quality of VOC blends, as was found for *Zea mays* (Gouinguené & Turlings, 2002). Within the present experiment, plants were not fertilised and all were grown in the same steamed soil but the pots may have received different amounts of water. Moreover, the PDMS approach is performed in a static headspace, in contrast to dynamic headspace sampling, in which a continuous air stream is applied to the collection unit (Tholl et al., 2006; Clancy et al., 2020). The application of a defined air stream may result in less variation in multi-laboratory comparisons. After VOC collection, we stored the PDMS tubes at −20 °C until further analysis as recommended by Kallenbach et al. (2014) since storage at room temperature is sufficient only for short-term storage.

To standardise the induction of VOCs, JA was applied to the substrate with a known JA concentration instead of adding herbivores, which could have shown a heterogeneous feeding pattern and thus introduce a further source of variation. Application of JA to the soil is likely more standardised as JA is then taken up with water and nutrients through the root system (van Dam et al., 2004). However, JA may also differently adsorbed by soil particles and not uniformly reach all roots. Moreover, JA was maybe not perfectly dissolved in all laboratories before application, introducing additional variation in the extent of induction. Applying JA directly on roots or shoots may provide a more natural herbivory simulation. Thereby, the type of JA application, i.e. either on roots or on shoots, shapes the induction of non-volatile leaf chemistry, as shown, for example, in *Arabidopsis thaliana* (Brassicaceae) (van Dam & Oomen, 2008), but also volatile leaf chemistry, as shown, for example, in *Brassica oleracea* (van Dam et al., 2010).

The organobromide 1-bromodecane is commonly used as internal standard in GC-MS measurements of VOCs, because it most likely does not occur in biological samples but has a similar behaviour as many terpenoids. This standard has already been applied e.g. in closed-loop stripping approaches (Quintana-Rodriguez et al., 2015) and automated solid-phase microextraction (Muchlinski et al., 2019). However, introducing 1-bromodecane as internal standard to the PDMS sampling protocol in the present experiment did not yield reproducible results. The compound had been applied on filter paper, which was inserted into the cup afterwards. Potential differences in evaporation time before adding the paper to the cup might have introduced a substantial variation in recovery both in terms of samples in which the internal standard could be detected and in amounts. Already during shipping to the participating laboratories, the solvent in which 1-bromodecane was solved, *n*-heptane, may have evaporated to different degrees. Whether applying 1-bromodecane directly onto PDMS tubes that are already in the cup for VOC collection might yield reproducible results needs to be examined. In addition, the 1-bromodecane normalised peak area varied across recipient laboratories, especially for the mono-chemotype, pointing to substantial variation in the sensitivity of instruments and instrument settings applied.

### 4.2 Variation due to analytical steps

In the present experiment, the VOC profiles from the cross-exchanged samples were analysed in the recipient laboratories utilising the respective available TD-GC-MS platforms. The high variation in number and quantities of VOCs detected in the different laboratories might have three reasons: (a) different analytical hardware, (b) variation in the settings of the TD-GC-MS measurements and (c) the experience with subsequent peak annotation.

Regarding (a): Analytical hardware differs in sensitivity as was, for example, found for LC-MS (Martin et al., 2015) but also for GC-MS (Lisec et al., 2006). Also, in the present study, the age of the used columns were not scored, but column aging influences the recovery and concentration of detected VOCs (Sangster et al., 2006). Regarding (b): Although we had standardised the measurement protocols as much as possible, some parameters were instrument-or laboratory-specific, e.g. the thermodesorption parameters, the type and length of the column, the injection temperature and the pressure as well as the flow of the carrier gas in the GC, and the scan speed in the MS. Many of these parameters have a strong influence on compound detection (Chow et al., 2007; Even et al., 2021; Ho et al., 2011; Mametov et al., 2021). Regarding (c): Different experiences with peak annotation and integration of peaks can introduce further variation in analytical steps (Izumi et al., 2019). Although we exchanged a list of expected VOCs and retention time indices known for the tested *T. vulgare* chemotypes, the overlap of commonly detected peaks was surprisingly low and there was some variation in the RIs. Moreover, quality assurance & quality control (QA/QC) measures to obtain reproducible results are not uniform across metabolomics laboratories (Dunn et al., 2017), and should be considered particularly in collaborative research.

One drawback of our analyses was that in two recipient laboratories (L2 and L5) a solvent cut-off of the first minutes (until minute 7.0 and 9.8, respectively) was applied in the TD-GC-MS measurements. This is usually applied in liquid injection to avoid overloading of the MS with solvent but not needed in solvent-free headspace sampling (Dettmer et al., 2007; Kallenbach et al., 2015). Several green leaf volatiles and monoterpenoids already elute prior to the applied cut-off and could thus not be detected in these laboratories. The cut-off unfortunately also prevented the detection of santolina triene, which was among the characteristic VOCs, for the mixed-chemotype. Comparative analyses thus had to be restricted to a small set of VOCs that were measured and detected in all recipient laboratories.

### 4.3 Variation caused by post-analytical steps

With our ring trial, we aimed at assessing the recovery of VOC profiles of both *T. vulgare* chemotypes and changes in VOC profiles due to JA treatment. Using the PDMS approach, we found chemotype-specific variation in VOCs (focusing on the six common terpenoids). Thus, at least within the laboratories, the two chemotypes could be discriminated. The high variation introduced by both donor and recipient laboratory in the number of peaks per sample and the composition of the six common terpenoids indicates that both pre-analytical steps (mirrored in significant variation between donor laboratories) and analytical as well as post-analytical steps (mirrored in significant variation between recipient laboratories) contributed to the variation in this ring trial study. Alternative adsorbents such as polyacrylate or divinylbenzene (Jalili et al., 2020; Souza Silva et al., 2013) or combinations of tubes of distinct material may be used to test for their affinity towards different VOCs, which may reduce variation in analytical and post-analytical steps.

Finally, the JA treatment induced VOCs in a bi-directional manner across donor laboratories, which could also be due to variation already present in plants during pre-analytical steps of the study. Interestingly, while JA treatment is usually expected to lead to an increased release of VOCs (El-Wakeil et al., 2010; van Schie et al., 2007), we found no consistent induction pattern in the six VOCs detected across all donor laboratories. Induction of VOCs has been found to be quite complex (Baldwin et al., 2002; Hogenhout & Bos, 2011). In the present study, the JA treatment into the soil might not have been sufficiently effective to induce a reproducible response, particularly, as there were some difficulties in dissolving JA sufficiently in all participating laboratories.

### 4.4 Conclusion

Sampling plant-emitted VOCs provides insights into how plants may communicate with their environment (Dicke & Baldwin, 2010; Pierik et al., 2014; Raguso, 2008). The PDMS approach was invented to provide a robust, cheap and simple technique to passively sample VOCs at trace levels and serve as a more flexible alternative to the classical dynamic headspace sampling techniques, such as closed-loop stripping or push-and-pull systems (Kallenbach et al., 2014; Tholl et al., 2006). It eliminates the need for solvents, but VOC concentrations can only be semi-quantified (Tholl et al., 2006). Thus, a combination of VOC sampling approaches may be more suitable. Applying a ring trial, we reached only a limited extent of reproducibility, although our VOC collection and analytical approach was capable to detect chemotype-specific variation. Plants that *per se* exhibit such a high chemodiversity in their VOC profiles as *T. vulgare*, as well as a high sensitivity to touching and mechanical disturbances of the extracellular terpenoid-containing glands, might be particularly challenging for inter-laboratory comparisons. We can attribute some of the irreproducibility to problems of protocol implementation. Thus, we like to emphasise the importance of training experiments to reveal reproducible results in collaborative projects.

## Acknowledgements

We thank Stephanie Champion, Lukas Brokate, Paul Krahmer and Dustin Raeke for help with setting up the experiment.

## Author contributions

EJE, RJ, JPS, AS, SBU, NMvD and CM designed the study. Particularly EJE, with the help of RJ, TB and CM (L5), prepared the plants and further material, which was sent to the other laboratories and coordinated the ring trial. RAA, KSA, MP, RS, FS, and DZ collected the VOC samples and measured plant and climate variables. TB, RAA, MP, RS, SAYV and DZ performed the TD-GC-MS measurements. SE analysed the data and wrote the first draft of the manuscript together with CM, with input from EJE and RJ. All authors contributed with comments and suggestions.

## Declarations

### Ethical approval

Not applicable.

### Consent to participate

Not applicable.

### Consent for publication

All authors approved the manuscript for publication.

### Open access

Raw data will be provided via the Zenodo repository. The R code generated during statistical analyses will be provided via the GitHub repository of SE.

