## Supplement for "Inter-laboratory comparison of plant volatile analyses in the light of intra-specific chemodiversity"

### Inter-laboratory comparison of volatile organic compound analysis in the light of intra-specific chemodiversity

| **Table of contents** | **Page** |
|---|---|
| **Method S1:** PDMS sampling protocol | 2 |
| **Table S1:** Environmental conditions for growing of experimental plants | 6 |
| **Table S2:** Annotated VOCs in participating recipient laboratories | 7 |
| **Table S3:** Model coefficients from type III two-way analysis of variance with plant height and fresh weight of the sampled leaf as response factors | 9 |
| **Table S4:** Model coefficients from type II two-way analysis of variance with the number of leaves and fresh weight of shoots as response factors | 10 |
| **Table S5:** Model coefficients from permutational two-way analysis of variance on recovery of 1-bromodecane | 11 |
| **Table S6:** Model coefficients from two-way linear mixed-effects models (LMM) on number of detected peaks per donor laboratory. | 12 |
| **Table S7:** Model coefficients from two-way linear mixed-effects models (LMM) on number of detected peaks per recipient laboratory. | 13 |
| **Table S8:** Model coefficients from linear mixed-effects models (LMM) on $\log_2$ fold changes from JA treatment | 14 |
| **Figure S1:** Images illustrating the VOC sampling procedure | 15 |
| **Figure S2:** Measurements of VOC standards for comparison of response factors | 16 |
| **Figure S3:** Amount of 1-bromodecane and VOC background profiles detected in blank samples | 17 |
| **Supplementary references** | 18 |

#### Methods

##### Method S1: PDMS sampling protocol.

##### Preparation of PDMS tubes

PDMS tubes (1 mm internal diameter, 1.8 mm external diameter; Carl Roth, Karlsruhe, Germany) were cut into 5 mm long pieces with a standardised cutting device as in Kallenbach et al. (2015). For cleaning, the tubes were soaked two times in a 4:1 (v:v) acetonitrile:methanol solvent-mix, first for 3 h at 80 °C and second overnight at room temperature. After the solvent had fully evaporated, tubes were conditioned in the TD-GC at 230°C and a flow of 60 mL $min^{-1}$ for 30 min following Kallenbach et al. (2014).

##### Prerequisites for VOC collection

- Two plants were chosen from a stock of greenhouse plants at one of the donor laboratories (L5) that were initially grown from seeds collected near Bielefeld (mono-chemotype: 51°58.635 N, 51°58.635 E, mixed-chemotype: 51°59.031 N, 51°59.031E). One plant was a β-thujone mono-chemotype and the other belonging to a myroxide-santolina triene-artemisyl acetate mixed-chemotype.
- L5 provided plants from root cuttings, pots and steamed substrate from each of both chemotypes and VOC collection material to the other donor laboratories (L1-L4). Of each chemotype, five plants were grown in individual pots and three further pots with substrate only served as blank samples. Plants were grown in a climate chamber (Supplementary Table S1) until VOC collection and watered well approximately thrice a week.
- Laboratory gloves were worn while preparing the volatile collection and strongly scented deodorants or creams were avoided to assure that VOC profiles were not contaminated with external volatiles.

##### Leaf preparation for VOC collection

1. From each plant, the youngest fully developed leaf was chosen for VOC collection.
2. For each plant, a balloon stick was cut to the appropriate height of the sampled leaf. A polyethylene terephthalate (PET) cup (Wimex, Náchod, Czech Republic) was horizontally aligned to the balloon stick to avoid that sampled leaves slipped out of

the cup and attached using two strips of adhesive tape. Additionally, cups for blank pots (without experimental plants but containing soil) were prepared in the same way.

3. Cups were cleaned with 70% ethanol and tissue paper. After evaporation of the ethanol (after approximately five minutes), cups were ready for VOC collection at the sampled leaf per experimental plant.

**VOC collection during control stage**

1. Four hours after the onset of the photoperiod (approximately 10:00 AM), each sampled leaf was enclosed in a separate cup through the hole in the bottom of the cup. The position of the leaf stalk was marked with twine where the cup ends.
2. Plants were allowed to recover from handling stress for about 24 hours (approximately 10:00 AM the following day). As preparation for the following day, two curved tweezers were cleaned with 70% ethanol and wrapped in aluminium foil.
3. On the following day, a 5 µL glass syringe was cleaned by drawing up heptane three times and discarding it. The cleaned syringe was used to apply twice 5 µL (= 10 µL in total) of 100 ng $\mu L^{-1}$ 1-bromodecane solution on a 1 $cm^2$ filter paper piece (pre-cut in a glass petri dish) as internal standard. Using the tweezer, the paper disc was gently placed onto the leaf in the cup.
4. Using the second cleaned tweezer, 12 clean polydimethylsiloxane (PDMS) tubes were inserted through the hole in the dome lid of the cup. PDMS tubes were placed at the same position in each of the cups without direct contact to the leaf, to each other to the filter paper with the internal standard. Additionally, 12 PDMS tubes were placed into each empty, closed cups serving as blank samples.
5. After six hours, PDMS tubes were gently removed from the cups by opening the dome lid and closing it again afterwards.
6. Between each harvest of PDMS tubes per plant, used tweezers were cleaned with 70% ethanol and, after evaporation of the ethanol, used to divide PDMS tubes into six labelled glass vials with two PDMS tubes per vial. During this procedure, contact between PDMS tubes and the leaf was avoided. Glass vials were sealed with PTFE

tape. Glass vials were cross-exchanged with the recipient laboratories and one vial was kept back as a backup. Glass vials were labelled with a unique labelling code.

**Jasmonic acid (JA)-treatment**

1. Three hours past onset of the photoperiod (approximately 9:00 AM), a JA solution (0.5 mg JA per 10 mL double-distilled water with 0.1% (w/v) Triton X-100) was prepared in a 250 mL laboratory glass bottle. The bottle was shaken vigorously for 10 seconds to ensure mixture.

7. Four hours after the onset of the photoperiod (approximately 10:00 AM), 10 mL of the JA solution was injected in three portions around the stem and into the soil of each pot using a plastic syringe without needle. Between each injection, syringes were changed or cleaned with double-ionised water. Again, two curved tweezers were cleaned with 70% ethanol and wrapped in aluminium foil as preparation for the next day.

**VOC collection after JA treatment**

1. VOC collection was conducted in the same way as during the control stage.
2. All glass vials with PDMS tubes were cross-exchanged between participating donor laboratories (turning them into recipient laboratories) for TD-GC-MS measurement.

**Measurement of phenotypic parameters**

1. Before the start of the preparation for VOC collection during control stage, plant height [cm] and the number of fully expanded leaves was noted.
2. After VOC collection post JA treatment, the sampled leaf was cut at the position of the twine after the cup was carefully removed and the fresh weight [g] was determined. Additionally, the whole aboveground biomass was cut and the fresh weight [g] was determined.

**Delivery of material**

Material was delivered in four batches to each donor laboratory.

**First delivery:** PET cups (Wimex, Náchod, Czech Republic) with hole at the bottom, balloon sticks, filter paper discs for 1-bromodecane application (in glass Petri dishes), labelled 1.5 mL glass vials for PDMS tube storage, 250 mL glass bottles and plastic syringes for JA treatment

**Second delivery:** Plants, soil substrate, pots

**Third delivery:** JA, Triton X-100, coloured twine, 2 mL Eppendorf tubes for leaf material, PDMS tubes (1 mm internal diameter, 1.8 mm external diameter; Carl Roth, Karlsruhe, Germany; cut in 5 mm long pieces and prepared similar to the description in Kallenbach et al. (2014), tape to attach balloon sticks to cups, 1-bromodecane (as internal standard), 5 µL syringe, envelopes to cross-exchange samples.

**Tables**

**Table S1:** Environmental conditions of experimental plants. For each climate chamber and greenhouse, respectively, temperature, light duration and intensity and humidity were determined approximately at the height of the blank cups and sampled leaf, respectively, at the sampling events before and after the treatment with jasmonic acid (JA).

| **Condition** | **L1** | **L2** | **L3** | **L4** | **L5** |
|---|---|---|---|---|---|
| **Control** | | | | | |
| Air temperature [°C] | 21.6 | 21.2 | 19.5-29.5 | 20.0 | 22.4 |
| Light duration (light [h]:dark [h]) | 16:8 | 16:8 | 16:8 | 16:8 | 16:8 |
| Photosynthetic active radiation (PAR) [µmol photons $m^{-2}$ $s^{-1}$] | 202 | 250 | ~250 | 165 | 346 |
| Air humidity [rel. %] | 58 | 67 | 22-49 | 70 | 69 |
| **JA treatment** | | | | | |
| Air temperature [°C] | 21.6 | 21.2 | 19.5-29.5 | 20.0 | 22.2 |
| Light duration (light [h]:dark [h]) | 16:8 | 16:8 | 16:8 | 16:8 | 16:8 |
| Photosynthetic active radiation (PAR) [µmol photons $m^{-2}$ $s^{-1}$] | 197 | 250 | ~250 | 165 | 347 |
| Air humidity [rel. %] | 58 | 67 | 22-49 | 70 | 60 |

**Table S2:** Annotated VOCs in participating recipient laboratories. Recipient laboratories (L1, L2, L4, L5) annotated peaks based on extracted ion chromatograms (EIC) and peak spectrum matching based on laboratory-specific and external spectrum libraries (NIST, FFNSC). The retention index (RI) was obtained via co-measured alkane series (C7-C40). VOCs are ordered according to their retention index (mean of samples per recipient laboratory) in ascending order. Only VOCs are listed that eluted with an RI higher than 933 due to the cut-off that was applied for earlier eluting compounds in two recipient laboratories.

| VOC | L1 | L2 | L4 | L5 |
|---|---|---|---|---|
| 1-bromodecane (internal standard) | 1353.8 | 1361.4 | 1352.3 | 1355.5 |
| α-pinene | 935.5 | 942.6 | 935.6 | 923.2 |
| RI948_unknown | 947.5 | - | - | - |
| RI960_unknown | - | 961.1 | - | - |
| *trans*-pinane | 971.0 | - | - | - |
| *m*-menthane | 976.5 | - | - | - |
| *cis*-pinane | 981.5 | - | - | - |
| RI975_unknown | - | 981.3 | - | - |
| RI985_unknown | 983.6 | - | - | - |
| 6-methyl-5-hepten-2-one | - | 988.8 | - | - |
| RI989_unknown | - | 988.9 | - | - |
| decane | - | 1001.7 | - | - |
| *2*-methyldecane | - | 1009.1 | - | - |
| RI1011_unknown | 1011.5 | - | - | - |
| RI1019_unknown | - | 1018.6 | - | - |
| 2-ethylhexanol | - | - | - | 1029.3 |
| RI1030_unknown | - | 1031.8 | - | - |
| salicyl aldehyde | - | - | 1045.2 | - |
| artemisia ketone | - | - | 1063.6 | - |
| RI1066_unknown | - | 1064.6 | - | - |
| *cis*-sabinene hydrate | - | - | - | 1074.9 |
| artemisia alcohol | - | - | 1093.3 | - |
| tetrahydrolinalool | - | - | 1097.5 | - |
| nonanal | - | 1107.9 | - | - |
| RI1118_unknown | - | 1119. 0 | - | - |
| (*Z*)-myroxide | - | - | - | 1141.7 |
| artemisyl acetate | - | - | 1175.0 | - |
| RI1176_unknown | 1176.4 | - | - | - |
| *trans*-linalool ethyl | - | - | 1195.0 | - |
| dodecane | - | 1200.4 | - | - |
| α-terpineol | - | - | - | 1201.4 |
| decanal | - | 1211.2 | - | - |
| RI1294_unknown | 1294.5 | - | - | - |
| RI1365_unknown | - | 1366.9 | - | - |
| RI1374_unknown | - | 1375.9 | - | - |
| RI1386_unknown | - | 1386.6 | - | - |
| RI1404_unknown | - | 1399.9 | - | - |
| RI1414_unknown | - | 1418.5 | - | - |

| **VOC (continued)** | **L1** | **L2** | **L4** | **L5** |
|---|---|---|---|---|
| RI1424_unknown | 1425.0 | - | - | - |
| RI1427_unknown | - | - | - | 1426.5 |
| RI1432_unknown | 1428.8 | - | - | - |
| RI1430_unknown | - | - | 1429.9 | - |
| RI1440_unknown | 1438.8 | - | - | - |
| *trans*-geranylacetone | 1455.5 | - | - | - |
| RI1504_unknown | - | 1506.5 | - | - |
| RI1514_unknown | - | 1517.9 | - | - |
| RI1520_unknown | - | 1519.5 | - | - |
| RI1536_unknown | - | 1538.8 | - | - |
| RI1628_unknown | 1625.4 | - | - | - |
| hexyl salicylate | - | - | - | 1689.0 |
| β-pinene | 979.0 | 981.6 | - | - |
| myrcene | 992.0 | 989.0 | - | - |
| *trans*-sabinene hydrate | - | 1078.8 | 1068.6 | - |
| acetophenone | 1070.5 | 1082.8 | - | - |
| menthol | - | - | 1175.5 | 1183.5 |
| *cis*-ethyl linalool | 1190.1 | - | 1182.2 | - |
| camphene | 951.0 | 962.5 | - | 943.4 |
| *trans*-geranylacetone | - | - | 1454.0 | 1451.2 |
| sabinene | - | 981.5 | 979.0±0 | 970.2 |
| *cis*-3-hexenyl acetate | 1008.0 | - | 1006.2 | 1004.4 |
| benzyl alcohol | 1038.0 | - | 1034.8 | 1038.3 |
| dihydromyrcenol | 1073.0 | - | 1072.5 | 1073.2 |
| methyl salicylate | - | 1211.5 | 1201.3 | 1200.4 |
| *p*-cymene | 1027.0 | 1034.9 | 1026.1 | 1027.0 |
| limonene | 1031.0 | 1039.3 | 1030.9 | 1031.5 |
| eucalyptol | 1034.3 | 1043.6 | 1034.5 | 1035.4 |
| β-caryophyllene | - | 1448.9 | 1426.4 | 1435.0 |
| α-thujone | 1109.5 | 1118.9 | 1110.6 | 1111.7 |
| β-thujone | 1120.5 | 1130.2 | 1120.0 | 1123.5 |
| α-farnesene | 1509.0 | 1547.0 | - | 1506.5 |
| camphor | 1151.5 | 1165.7 | 1150.7 | 1156.6 |
| borneol | 1173.2 | 1189.7 | 1171.8 | 1180.5 |

**Table S3:** Model coefficients from type III two-way analysis of variance (ANOVA) with plant height and fresh weight of the sampled leaf as response factors. Response variables were transformed to facilitate normality. Only the interaction of the main factors donor laboratory (D) and chemotype (C) is shown. Total sample size *n* = 50. Abbreviations: BC – Box-Cox transformation; SumSq – sum of squares, Df – degree of freedom.

| | **Plant height [cm]** | **FW leaf [g]** |
|---|---|---|
| *Sample size* | 50 | 50 |
| *Transformation* | - | BC |
| $SumSq_{Residuals}$ | 164 | 24.51 |
| $Df_{Residuals}$ | 40 | 40 |
| *Intercept* | | |
| *Sum of squares* | 14.67 | 0.00 |
| *Df* | 1 | 1 |
| *F-value* | 3576.79 | 0.00 |
| *p-value* | < 0.001 | 1.00 |
| *Interaction D x C* | | |
| *SumSq* | 89 | 7.33 |
| *Df* | 4 | 4 |
| *F-value* | 5.42 | 2.99 |
| *p-value* | 0.001 | 0.030 |

**Table S4:** Model coefficients from type II two-way analysis of variance with the number of leaves and fresh weight of shoots as response factors. Response variables were transformed to facilitate normality. The interaction of the main factors donor laboratory (D) and chemotype (C) was dropped to facilitate model fit. Total sample size *n* = 50. Abbreviations: OQN – ordered-quantile normalisation; SumSq – sum of squares, Df – degree of freedom.

| | **Number of leaves** | **FW shoots [g]** |
|---|---|---|
| *Transformation* | OQN | - |
| $SumSq_{Residuals}$ | 19.89 | 50.52 |
| $Df_{Residuals}$ | 44 | 44 |
| *Intercept* | - | - |
| *Donor laboratory (D)* | | |
| *SumSq* | 25.22 | 88.01 |
| *Df* | 4 | 4 |
| *F-value* | 13.95 | 19.16 |
| *p-value* | < 0.001 | < 0.001 |
| *Chemotype (C)* | | |
| *SumSq* | 0.47 | 6.16 |
| *Df* | 1 | 1 |
| *F-value* | 1.03 | 5.63 |
| *p-value* | 0.315 | 0.025 |

**Table S5:** Model coefficients from permutational two-way analysis of variance ($LM_{Perm}$) on recovery of 1-bromodecane. The number of samples per recipient laboratory, in which the internal standard was detected, was used as response variable. The non-significant interaction between the explanatory variables donor laboratory and chemotype was removed in the final model. Abbreviations: $SumSq_R$ – permutational sum of squares; $MeanSq_R$ – permutational mean squares; Iter – iterations.

| **Explanatory variable** | **Df** | **$SumSq_R$** | **$MeanSq_R$** | **Iter** | ***p*-value** |
|---|---|---|---|---|---|
| Donor laboratory | 4 | 0.39 | 0.10 | 1775 | 0.120 |
| Chemotype | 1 | 0.03 | 0.03 | 238 | 0.298 |
| Residuals | 32 | 1.65 | 0.05 | | |

**Table S6:** Model coefficients from two-way linear mixed-effects models (LMM) on number of detected peaks per donor laboratory. The counts, i.e. the number of detected peaks, was used as response variable. Donor laboratory, chemotype and their interaction were applied as fixed factors. Treatment, recipient laboratory as well as plant individual nested within recipient laboratory were applied as random factors. The non-significant interaction between the fixed factors was removed in the final model. Abbreviations: Df – Satterthwaite's degrees of freedom (Num – Numerator; Den – Denominator); SumSq – sum of squares; MeanSq –mean squares.

| **Fixed factors** | **SumSq** | **MeanSq** | **NumDf** | **DenDf** | ***F*-value** | ***p*-value** |
|---|---|---|---|---|---|---|
| *Donor laboratory (D)* | 626.24 | 156.56 | 4 | 375.01 | 18.38 | < 0.001 |
| *Chemotype (C)* | 2.21 | 2.21 | 1 | 375.00 | 0.26 | 0.611 |

**Table S7:** Model coefficients from two-way linear mixed-effects models (LMM) on number of detected peaks per recipient laboratory. The counts, i.e. the number of detected peaks, was used as response variable. Recipient laboratory, chemotype and their interaction were applied as fixed factors. Treatment, Recipient laboratory as well as plant individual nested within recipient laboratory were applied as random factors. The non-significant interaction between the fixed factors was removed in the final model. Abbreviations: Df – Satterthwaite's degrees of freedom (Num – Numerator; Den – Denominator); SumSq – sum of squares; MeanSq – mean squares.

| **Fixed factor** | **SumSq** | **MeanSq** | **NumDf** | **DenDf** | ***F*-value** | ***p*-value** |
|---|---|---|---|---|---|---|
| *Donor laboratory (D)* | 2667.30 | 8879.10 | 3 | 374.67 | 1039.46 | < 0.001 |
| *Chemotype (C)* | 2.20 | 2.20 | 1 | 373.84 | 0.26 | 0.611 |

**Table S8:** Model coefficients from two-way linear mixed-effects models (LMM) on $log_2$ fold changes from JA treatment. LMMs were conducted separately for each chemotype. Donor laboratory (D), compound (Com) and their interaction (D x Com) were applied as fixed factors. Recipient laboratory as well as plant ID nested within recipient laboratory were applied as random factors. Abbreviations: Df – Satterthwaite's degrees of freedom (Num – Numerator; Den – Denominator); SumSq - sum of squares; MeanSq –mean squares.

| **Fixed factor** | **SumSq** | **MeanSq** | **NumDF** | **DenDf** | ***F*-value** | ***p*-value** |
|---|---|---|---|---|---|---|
| *Mono-chemotype* | | | | | | |
| *Donor laboratory (D)* | 59.03 | 14.76 | 4 | 554.99 | 18.76 | < 0.001 |
| *Compound (Com* | 13.95 | 2.79 | 5 | 550.21 | 3.55 | 0.004 |
| *Interaction D x Com* | 29.65 | 1.48 | 20 | 550.21 | 1.88 | 0.012 |
| *Mixed-chemotype* | | | | | | |
| *Donor laboratory (D)* | 11.38 | 2.84 | 4 | 555.14 | 3.22 | 0.012 |
| *Compound (Com* | 8.66 | 1.73 | 5 | 550.05 | 1.96 | 0.082 |
| *Interaction D x Com* | 29.70 | 1.48 | 20 | 550.05 | 1.68 | 0.032 |

**Figures**

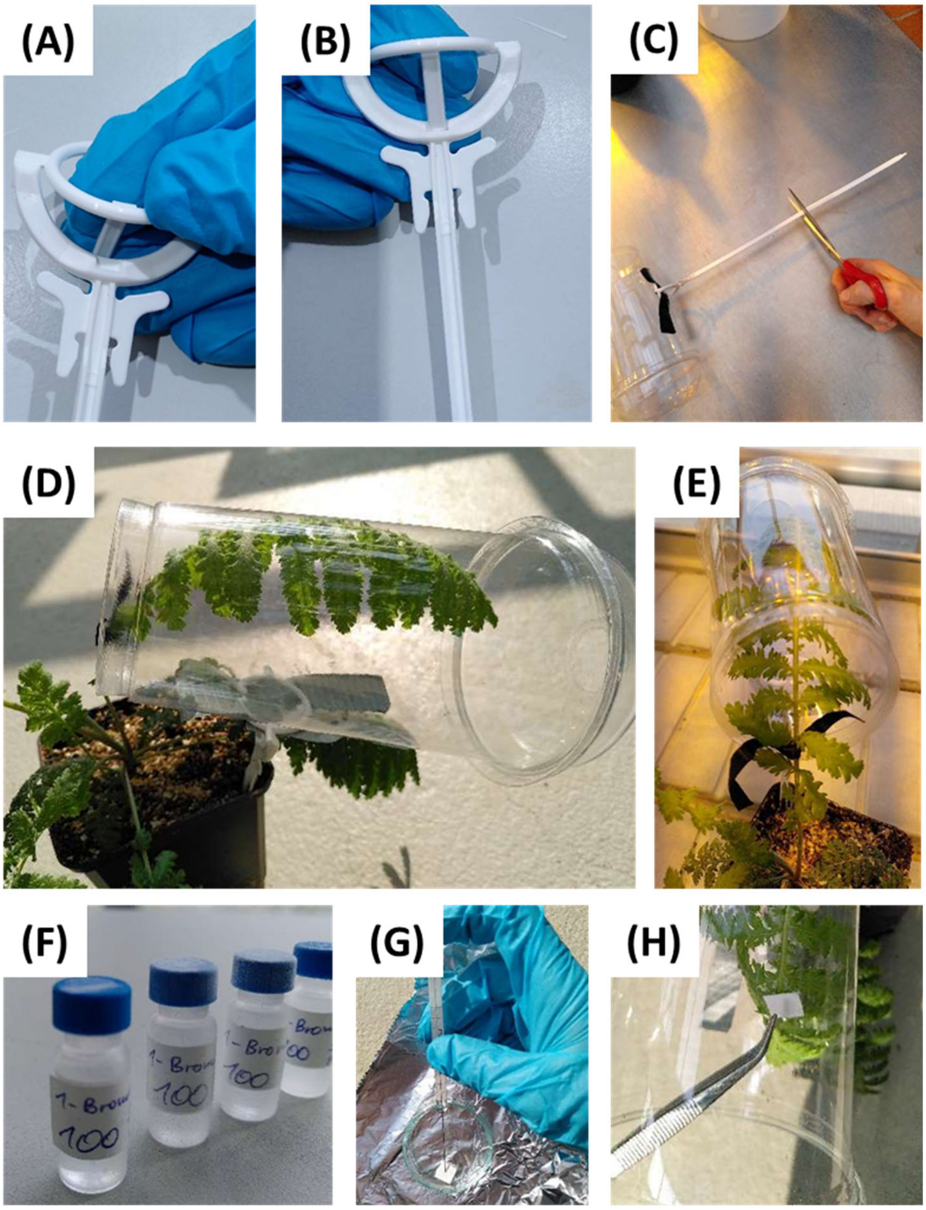


**Figure S1:** Images illustrating the VOC sampling procedure. Young and intact leaves of *Tanacetum vulgare* were enclosed in **(A-C)** cups fixed with balloon sticks, **(D-E)** PDMS tubes were carefully placed onto the enclosed leaf and **(F-H)** 1-bromodecane was applied as internal standard on filter paper and placed into the cup next to the leaf.

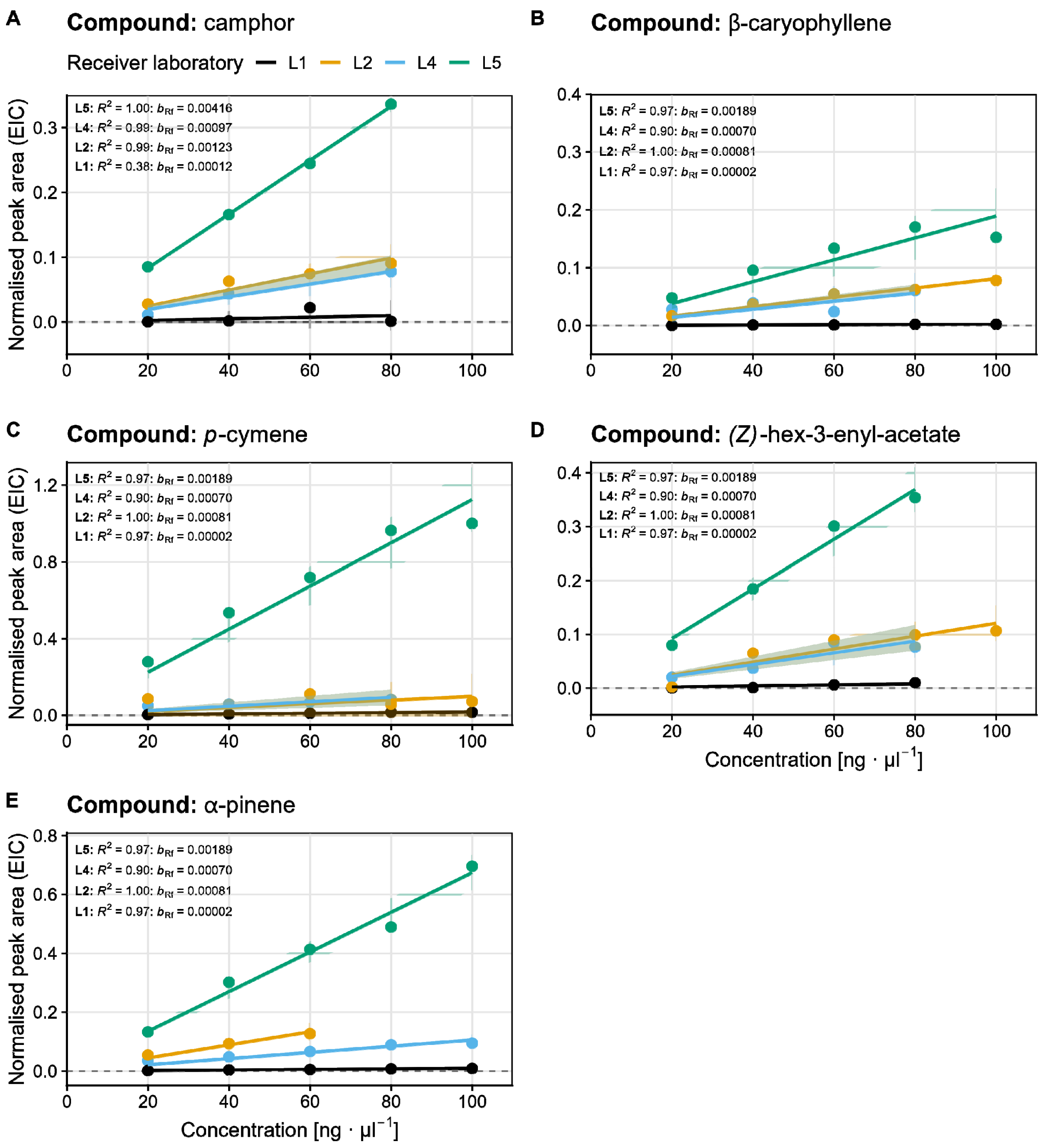


**Figure S2:** Measurements of VOC standards for comparison of response factors. A minimum of three concentration levels along a dilution series (20 ng µL$^{-1}$, 40 ng µL$^{-1}$, 60 ng µL$^{-1}$, 80 ng µL$^{-1}$, 100 ng µL$^{-1}$) was measured for **(A)** camphor, **(B)** β-caryophyllene, **(C)** *p*-cymene, **(D)** (*Z*)-hex-3-enyl-acetate, and **(E)** α-pinene under laboratory-specific TD-GC-MS settings of the recipient laboratories (L1, L2, L4, L5). Response factors were defined as the zero-intercept slope of the linear regression line of peak area against concentration per laboratory-specific measurement. For readability, the y-axis has been rescaled between zero and one (normalised peak area based on EIC), and upper limits of subfigures adjusted according to the compound-specific maximum value. Abbreviations: $R^2$ – coefficient of determination; $b_{Rf}$ – slope of the linear regression line.

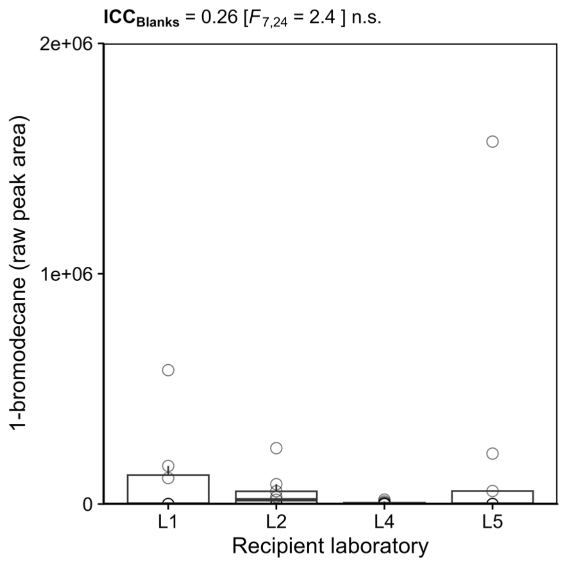


**Figure S3:** Amount of 1-bromodecane detected in blank samples. The reproducibility of 1-bromodecane recovery was analysed as the intra-class correlation coefficient (ICC) based on raw peak areas visualised with box-and-whisker plots (median within 50 % of data in boxes and 1.5-times the inter-quartile range as whiskers). One very high value (5.3e+06) is not shown for readability. Abbreviations: Asterisks denote significance levels in ICC analysis; n.s. – not significant.
